# The relationship between response dynamics and the formation of confidence varies across the lifespan

**DOI:** 10.1101/2022.06.14.493033

**Authors:** Helen Overhoff, Yiu Hong Ko, Gereon R. Fink, Jutta Stahl, Peter H. Weiss, Stefan Bode, Eva Niessen

**Author notes:** Corresponding author. Cognitive Neuroscience, Institute of Neuroscience and Medicine (INM-3), Research Centre Jülich, Leo-Brandt-Str. 5, 52425 Juelich, Germany., *Email:.

## Abstract

Accurate metacognitive judgements, such as forming a confidence judgement, are crucial for goaldirected behaviour but decline with older age. Besides changes in the sensory processing of stimulus features, there might also be changes in the motoric aspects of giving responses that account for age-related changes in confidence. In order to assess the association between confidence and response parameters across the adult lifespan, we measured response times and peak forces in a four-choice flanker task with subsequent confidence judgements. In 65 healthy adults from 20 to 76 years of age, we showed divergent associations of each measure with confidence, depending on decision accuracy. Participants indicated higher confidence after faster responses in correct but not incorrect trials. They also indicated higher confidence after less forceful responses in errors but not in correct trials. Notably, these associations were age-dependent as the relationship between confidence and response time was more pronounced in older participants, while the relationship between confidence and response force decayed with age. Our results add to the notion that confidence is related to response parameters and demonstrate noteworthy changes in the observed associations across the adult lifespan. These changes potentially constitute an expression of general age-related deficits in performance monitoring or, alternatively, index a failing mechanism in the computation of confidence in older adults.

## 1. Introduction

Humans can report a subjective sense of confidence that is closely related to the accuracy of their actions. This ongoing monitoring of decisions and their execution is called metacognition and includes the evaluation of behaviour and the detection of occurring errors (Fleming & Dolan, 2012). Accurate metacognitive judgements should lead to adaptive behaviour adjustments and are thus crucial for all activities. Undetected errors (i.e., incorrect metacognitive judgements) might have severe implications for real-life scenarios because they may not trigger the required adjustments for future actions and decisions (Wessel et al., 2018).

### 1.1. Age-related decline in metacognitive accuracy

Research on metacognitive performance across the adult lifespan has consistently pointed towards a decline in older age. When participants were asked to report committed errors in an easy choice-reaction task, the detection rates declined with age, even when task performance was comparable (Harty et al., 2017; Niessen et al., 2017). In our previous publication, using the same dataset described in this study (Overhoff et al., 2021), we asked participants to rate their confidence after each decision on a four-point scale, and we concordantly revealed a decline in metacognitive performance across the lifespan. The accuracy of these ratings decreased gradually with higher age, reflecting that older adults were less aware of their errors and rated correct responses with lower confidence compared to younger adults (see also Palmer et al., 2014). As of today, the question of which factors are related to this selective decline remains open.

### 1.2. Computation of confidence

In order to understand the age-related decline in metacognitive performance, it is essential to understand the basic mechanisms underlying the computation of confidence. It is still unclear which information is used to compute confidence, and how input from different sources is weighted (Charles and Yeung, 2019; Feuerriegel et al., 2021). For instance, confidence has been related to the strength of stimulus evidence, stimulus discriminability (Charles and Yeung, 2019; Turner et al., 2021b; Yeung and Summerfield, 2012), or instructed time pressure (Vickers and Packer, 1982). Furthermore, growing evidence suggests that the interoceptive feedback of a motor action while giving a response might be another source of information contributing to the formation of confidence about the decision (Fleming et al., 2015; Gajdos et al., 2019; Kiani et al., 2014; Palser et al., 2018; Siedlecka et al., 2021; Turner et al., 2021a). Fleming and colleagues (2015) investigated the interaction of confidence and motor-related activity by delivering single-pulse transcranial magnetic stimulation (TMS) to the dorsal premotor cortex. This perturbation did not affect task performance, but crucially, it did affect the accuracy of subsequent confidence judgements, i.e., the degree to which the judgements matched the observed performance. This finding indicates that action-specific cortical activations might contribute to confidence. In line with this assumption, confidence ratings have been shown to be more accurate if the preceding decision required a motor action (Pereira et al., 2020; Siedlecka et al., 2021). For instance, Siedlecka and colleagues (2021) recently showed that metacognitive accuracy was higher after decisions requiring a key press than decisions which were indicated without a motor action. Taken together, these findings suggest that features of the motor response indicating a given decision might influence the confidence ratings about this decision. Therefore, further investigations of how confidence is reflected in different response parameters are warranted.

### 1.3. Differential relationship between confidence and response parameters

A response can be characterised by different dimensions. The most commonly used output variable is time, usually response time or movement time. A robust finding across studies is a negative relationship between response times for the initial decision and subsequent confidence ratings (Fleming et al., 2010; Kiani et al., 2014; Rahnev et al., 2020). Intuitively, one might assume that the degree of confidence is expressed in the time taken to make the decision, i.e., the less confident we are about a decision, the longer it should take to respond. However, another possible explanation is that the monitoring system uses the interoceptive signal of a movement produced by the response as an informative cue about the difficulty of the decision (Fleming and Daw, 2017; Kiani et al., 2014). Accordingly, if an easy decision led to a fast response, the internal read-out could boost subjective confidence. A recent study provided evidence for the directional effect of movement time (i.e., the time from lifting to dropping a marble) on confidence (Palser et al., 2018). In this study, movement speed was experimentally manipulated by instructing participants to move faster than they naturally would, and this manipulation resulted in declined metacognitive accuracy. Nevertheless, temporal parameters do not capture all aspects of a movement. For instance, subthreshold motor activity (i.e., partial responses) cannot be detected by classical RT recordings but rather by recording muscle activity. However, partial responses have also been shown to affect reported confidence (Ficarella et al., 2019; Gajdos et al., 2019). An informative motor parameter of a response is the applied force, which is often measured in its peak force, i.e., the maximum exerted force during a response action. Notably, peak force and response time index distinct processes as they show divergent behavioural patterns (i.e., small to no correlation) across experimental manipulations (Cohen and van Gaal, 2014; Franz and Miller, 2002; Stahl and Rammsayer, 2005).

Contrary to the well-known negative relationship between confidence and response time, the association between confidence and response force has rarely been investigated. Recently, Turner et al. (2021a) examined this relationship by explicitly manipulating the degree of physical effort that had to be exerted to give a response. When participants were prompted to submit their response to a perceptual decision with varying force levels, participants reported higher confidence in their decisions when their response peak force was higher. Notably, requiring participants to produce a specific (and comparably high) degree of force (as mandated in the experiment) is fundamentally different from measuring naturally occurring force patterns of a response (in terms of a dependent measure). The latter was done, for example, in a study by Bode and Stahl (2014), who found that naturally occurring peak force was lower in errors compared to correct responses. It was suggested that this might indicate a process in which low force in error trials signifies an unsuccessful attempt to stop the already initiated response, which requires early and fast error detection (Bode and Stahl, 2014; Ko et al., 2012; Stahl et al., 2020). However, error detection or confidence was not directly assessed, rendering comparison between these two studies difficult.

The relationship between different response parameters and confidence has not been systematically assessed in the context of healthy ageing. While response and movement times are slower and more variable with older age, findings of age-related changes in response force are inconsistent (Bunce et al., 2004; Dully et al., 2018; Salthouse, 2000). Some studies showed delayed and altered electrophysiological signatures of motor processing in older age (e.g., lateralised readiness potential (LRP)/ movement-related potential (MRP) and mu/ beta desynchronisation; Falkenstein et al., 2006; Quandt et al., 2016; Sailer et al., 2000). In contrast, electromyographic or force recordings of motor responses revealed similar patterns in younger and older adults (Dully et al., 2018; Falkenstein et al., 2006; Van Der Lubbe et al., 2002; Yordanova et al., 2004). Notably, these studies did not assess error awareness or confidence. Therefore, it is warranted to specifically examine the associations between confidence and response time and between confidence and response force and to investigate whether these associations change across the adult lifespan.

### 1.4. Objectives

The present study constitutes the first comprehensive assessment of the association between metacognitive accuracy and two main response parameters across the adult lifespan. We intended to answer the following questions: First, what are the relationships between decision confidence and response time on the one hand, and peak force of a response (as it naturally occurs, i.e., without specific instruction or experimental manipulation) on the other hand? Second, do these relationships between confidence and response parameters change with age? Additionally, we were interested in investigating the potential moderating effect of accuracy because many studies on decision confidence only assessed the relationship between a given response parameter and confidence in correct responses. However, we can only understand the computation of confidence when considering errors (Charles and Yeung, 2019; Dotan et al., 2018; Peters et al., 2017). Concerning response time, for instance, the well-known negative relationship with confidence is inverted for errors when the confidence rating is allowed to indicate error detection (i.e., a rating scale was used that ranged from certainty in being correct to certainty in being wrong; Pereira et al., 2020).

We expected significant associations between confidence judgements and parameters of the response (Fleming et al., 2015; Pereira et al., 2020; Rahnev et al., 2020; Turner et al., 2021b). In particular, response time was expected to decrease with higher confidence for correct trials (Dotan et al., 2018; Kiani et al., 2014; Rahnev et al., 2020) and to increase with higher confidence for errors (Pereira et al., 2020). We tentatively hypothesised a positive relationship between response force and confidence for errors and correct responses (Bode and Stahl, 2014; Ko et al., 2012; Turner et al., 2021a).

Most importantly, we intended to explore age-related changes in the associations between response parameters and confidence without having a priori hypotheses about the direction of possible effects due to a lack of previous studies on this topic. If we find divergent patterns across the lifespan, this might encourage research on the causal relationship between response parameters and confidence.

## 2. Methods

### 2.1. Participants

Eighty-two participants were recruited and received monetary compensation for their participation in the experiment. Data from seventeen participants had to be discarded due to: symptoms of depression (*N* = 1, Beck’s Depression Inventory score higher than 17; BDI; Hautzinger, 1991), poor behavioural performance (*N* = 8, more than 30% invalid trials, error rate higher than chance, here 25%), or a behavioural pattern that was indicative of an insufficient understanding or implementation of task demands (*N* = 8, inspection of individual datasets for a combination of errors in the colour discrimination test described below, near chance task performance, frequent invalid trials, and biased use of single response keys). This resulted in a final sample for analysis of sixty-five healthy, right-handed adults (age = 45.5 ±2.0 years [all results are indicated as mean ±standard error of the mean; *SEM*]; age range = 20 to 76 years; 26 female, 39 male) with (corrected to) normal visual accuracy, no colour-blindness, no signs of cognitive impairment (Mini-Mental-State Examination score higher than 26; MMSE; Folstein et al., 1975) and no history of psychiatric or neurological diseases.

The current study’s data has been used previously (Overhoff et al., 2021). The same exclusion criteria regarding the neuropsychological assessment and the task performance were applied, resulting in the same subsample included in the analyses. In the previous publication, we thoroughly examined the metacognitive performance and its relation to behavioural parameters (response accuracy, response time, behavioural adjustments) as well as two electrophysiological potentials (i.e., the error/correct negativity, N_e_, and the error/correct positivity, P_e_; for detailed results and discussion thereof, see Overhoff et al., 2021). We did not report or analyse any response force measures in the previous publication.

The experiment was approved by the ethics committee of the German Psychological Society (DGPs). All participants gave written informed consent, and the study followed the Declaration of Helsinki.

### 2.2. Stimuli

The experiment consisted of a colour version of the Flanker task (Eriksen and Eriksen, 1974) with four response options intended to increase conflict and thereby the number of errors while ensuring feasibility for participants of all ages. Four target colours were mapped onto both hands’ index and middle fingers. In each trial, we presented one central, coloured target square flanked by two squares on the left and right side, respectively. Participants had to respond to the central target by pressing the corresponding finger. The flankers were presented slightly before the target appeared to increase their distracting effect. Flankers could be of the same colour as the target (congruent condition), of one of three additional neutral colours that were not mapped to any response (neutral condition), or of another target colour (incongruent condition).

### 2.3. Experimental Paradigm

Each trial started with the presentation of a white fixation cross on black background for 500 ms. The fixation cross was replaced by the two flankers, followed by the target after 50 ms and the two flankes and the target remained on screen for another 100 ms. Participants pressed their left or right index or middle finger to indicate their decision (see Figure 1B). Participants were instructed to respond as fast and accurately as possible. A black screen was presented until a response was registered (max. 1200 ms) and an additional 800 ms before presenting the confidence rating. For this rating, participants indicated their confidence in the decision on a four-point scale comprising the options ‘surely wrong’, ‘maybe wrong’, ‘maybe correct’, and ‘surely correct’ (max. 2000 ms). A jittered intertrial interval of 400 to 600 ms preceded the subsequent trial. If no response was registered in the decision task, the participants received feedback about being too slow, and the trial was terminated. The sequence of an experimental trial is depicted in Figure 1A.

**Figure 1.**
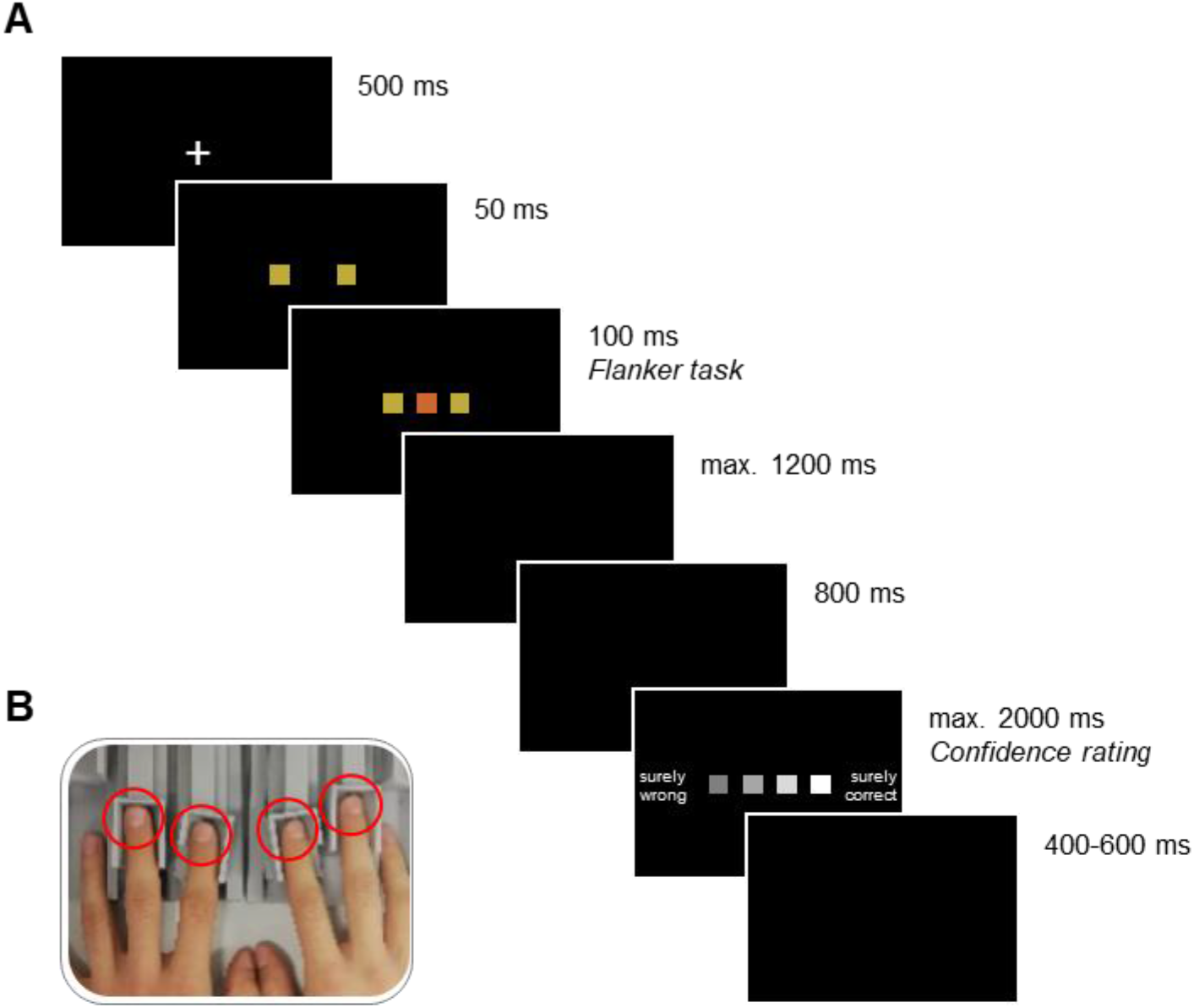
(A) Trial structure (here, incongruent condition illustrated). Each trial commenced with the presentation of a fixation cross. Subsequently, two coloured squares (flankers) were presented, and a third square (of the same or a different colour; target) was added shortly after. The stimuli disappeared after 100 ms and the ensuing black screen, where participants were instructed to make a response by pressing one of four response keys mapped onto one colour each, remained until a response was registered (maximum 1,200 ms). If no response was given, the German words for ‘too slow’ were shown, and the trial was terminated. Otherwise, after another black screen, the confidence rating scale was presented, which remained on the screen until a judgement (the four fingers were mapped onto the squares according to their spatial location) was made (maximum 2,000 ms). The next trial started after another black screen of random duration between 400 and 600 ms. (B) Force-sensitive response keys. Left and right index and middle fingers (red circles) were placed on adjustable finger rests.

### 2.4. Procedures

Prior to testing, we collected demographic details, and the participants conducted a brief colour discrimination test without any time pressure or cognitive load to ensure that they were capable of correctly discriminating the stimulus colours used in the experiment. The neuropsychological tests for assessing the exclusion criteria (BDI; MMSE; Edinburgh Handedness Inventory, EHI; Oldfield, 1971) were administered after the main experiment.

Participants first performed 18 practice trials without confidence rating, receiving feedback about their accuracy, which could be repeated if necessary. Two practice blocks of 72 trials without feedback followed. Another practice block then introduced the confidence rating. The actual experiment consisted of five blocks with 72 trials each, with optional breaks after each block. The electroencephalogram (EEG) was recorded throughout the testing session. Note that the EEG results have been reported in our previous publication (Overhoff et al., 2021).

### 2.5. Apparatus

The participants were seated in a noise-insulated and dimly lit testing booth at a viewing distance of 70 cm to the screen (LCD monitor, 60 Hz). A chin rest minimised non-task related movements.

For response recording, we used force sensitive keys with a sampling rate of 1024 Hz and a high temporal resolution that is superior to standard keyboards (Figure 1B; Stahl et al., 2020). The keys were calibrated to the fingers’ weight before and during the experiment. The keys could be adjusted to the hand size, and a comfortable hand position was ensured by a wrist rest. An applied force was registered as a response when it exceeded a threshold of 40 cN.

The colour discrimination test was programmed using Presentation software (Neurobehavioural Systems, version 14.5) and the main task using uVariotest software (version 1.978).

### 2.6. Analysis

Response time (RT) was defined as the time from target stimulus presentation to the initial crossing of the response force threshold of 40 cN by any response key. Peak force (PF) was defined as the maximum of a force pulse following the crossing of the threshold. Additionally, we measured the time from response onset to the time of the PF (only used for the exclusion of trials).

We excluded from the analysis: invalid trials, which were too slow, responses without confidence rating, responses with an RT below 200 ms (indicating premature responding), a PF below 40 cN (indicating incomplete or aborted responding), a time to PF of more than three standard deviations above the mean (indicating that the response was not of the expected ballistic nature), and recording artefacts (implausible time between response onset and time of the PF, incorrect identification of response key in case of multiple responses).

As a first step, to characterise the distribution of the behavioural parameters of interest independent of confidence, we computed paired samples *t*-tests at the group level to compare RT, PF, and their dispersion between correct and incorrect trials. For the investigation of age-related effects, we used a series of linear regressions with the predictor age for each of the following variables: error rates (ER; the proportion of valid responses that were incorrect), mean confidence ratings, mean RT and PF, and standard deviation of RT and PF. The latter analyses were performed separately for errors and correct responses.

Next, data were analysed using generalised linear mixed-effects models (GLMMs) with a beta distribution using the glmmTMB package (version 1.0.2.1; Brooks et al., 2017) in R (version 4.0.5; R Core Team, 2021). We chose this modelling approach because the beta distribution is assumed to better account for data that are not normally distributed and doubly bounded (i.e., having an upper and a lower bound; here: 1, “surely wrong”, and 4, “surely correct”), which applies to our confidence data (Verkuilen and Smithson, 2012). All continuous predictor variables were mean centred and scaled for model fitting, and confidence was scaled to the open interval (0,1; i.e., the range is slightly compressed to avoid boundary observations; Verkuilen and Smithson, 2012). Analyses were again conducted separately for correct responses and errors.

We examined the effects of age and the two parameters (RT, PF) of the response on confidence ratings using the following regression model structures (separately for the subsets of errors and correct responses):

1. Confidence ∼ Age + (RT | Participant)
2. Confidence ∼ RT*Age + (RT | Participant)
3. Confidence ∼ PF*Age + (RT | Participant)
4. Confidence ∼ RT*PF*Age + (RT | Participant)

RT and PF were used as fixed effects, and age was included as a covariate due to its documented negative effect on metacognitive accuracy (i.e., a negative effect on confidence for correct responses and a positive effect on confidence for errors; Overhoff et al., 2021; Palmer et al., 2014). For the most complex model, we considered an interaction term between all three factors, as RT and PF are known to vary across age (Dully et al., 2018), and previous work suggests potential interactions between RT and PF (Bode and Stahl, 2014; Gajdos et al., 2019). We fitted random intercepts for participants, allowing their mean confidence ratings to differ. If possible and the models converged, random slopes by participant were added for the predictors of interest to account for individual differences in the degree to which these were related to the confidence ratings (Barr et al., 2013). Models were checked for singularity and multicollinearity by calculating the variance inflation factor (VIF) using the performance package (version 0.7.2; Lüdecke et al., 2021).

We compared model fits including all effects of interest (model 4) to models including only one (models 2, 3) or no effect of interest (model 1) using likelihood ratio tests, and computed Wald *z*-tests to determine the significance of each coefficient. This means that, if a model including one predictor of interest (e.g., RT) fits the data better than a model including no effect of interest, this predictor has a relevant effect on confidence, and its inclusion in the model allows for a better prediction of participants’ ratings.

To follow up on significant interaction effects between age and the predictors of interest, we calculated slopes for three values of age (the mean and one standard deviation above and below the mean). Additionally, for statistical analysis of the transitions between these values, we computed Johnson-Neyman intervals using an adapted version of the *johnson_neyman* function of the interactions package (version 1.1.0; Long, 2019). This analysis reveals whether the statistical effect of the response parameters on confidence is conditional on the entire range of the moderator age, or just a sub-range, thus providing bounds for where the observed interaction effect is significant.

## 3. Results

### 3.1. Overview of response parameters

On average, participants had an error rate of 15.4 ± 1.6 %. Correct trials had a mean RT of 709.2 ± 11.5 ms and were faster [*t*(64) = −3.01, *p* = .004] and had a smaller standard deviation [*t*(64) = −5.53, *p* < .001] than error trials with an RT of 734.3 ± 13.9 ms. The mean peak force (PF) was higher for correct trials (236.2 ± 13.1 cN) compared to errors [191.7 ± 10.3 cN; *t*(64) = 5.51, *p* < .001] but did not differ in its standard deviation [*t*(64) = −0.21, *p* = .836; see supplementary Figure S1].

### 3.2. Effect of age on response parameters

We have already reported the relationship between age and error rate, RT, and confidence in our previous publication (Overhoff et al., 2021) based on a slightly different subset of trials to the one used here (due to additional force-related exclusions of trials in this study). Our initial results were confirmed using a series of linear regression analyses, each using age as the predictor for one of the following variables: We found that, at group level, the error rate increased with age [*F*(1,63) = 34.12, *p* < .001, β = 0.005, *SE* = 0.001, *t* = 5.84]. RT increased with age for correct [*F*(1,63) = 27.07, *p* < .001, β = 3.115, *SE* = 0.599, *t* = 5.20] and incorrect responses [*F*(1,63) = 10.16, *p* = .002, β = 2.568, *SE* = 0.806, *t* = 3.19], while age did not significantly predict PF for either type of response [correct: *F*(1,63) = 0.02, *p* = .884, β = 0.120, *SE* = 0.816, *t* = 0.15; error: *F*(1,63) = 1.33, *p* = .254, β = 0.734, *SE* = 0.637, *t* = 1.15]. RTs were more variable with higher age for correct responses [*F*(1,63) = 7.43, *p* = .008, β = 0.477, *SE* = 0.175, *t* = 2.73], but not errors [*F*(1,63) = 0.04, *p* = .837, β = 0.053, *SE* = 0.255, *t* = 0.21]. Similar to the mean PF, the standard deviation of PF did not change with age [correct: *F*(1,63) = 0.15, *p* = .704, β = 0.164, *SE* = 0.429, *t* = 0.38; error: *F*(1,63) = 0.08, *p* = .774, β = 0.137, *SE* = 0.475, *t* = 0.30]. These results are illustrated in the supplementary Figure S1.

The mean confidence (in the decision being correct, on a scale from 1 to 4) for correct responses (3.82 ± 0.02 for the entire sample) decreased with age [*F*(1,63) = 22.42, *p* < .001, β = − 0.007, *SE* = 0.002, *t* = −4.74]. This finding indicates that the older participants were, the less confident they were in being correct when responding correctly. Contrarily, the mean confidence for errors (2.35 ± 0.08 for the entire sample) increased with age [*F*(1,63) = 21.96, *p* < .001, β = 0.019, *SE* = 0.004, *t* = 4.69; see supplementary Figure S2]. Hence, the older the participants were, the less sure they were that the decision was wrong when making an error. We have recently described this phenomenon as an age-related tendency to use the middle of the confidence scale, pointing towards increased uncertainty in older adults (Overhoff et al., 2021).

### 3.3. Modelling of confidence

Variance inflation factors across all models with interactions were < 2.03, indicating low collinearity (< 5; James et al., 2013) between the predictors, and the models were not overfitted, as the fits proved not to be singular.

#### 3.3.1. Confidence in correct decisions

We computed likelihood ratio tests to compare the model fit of the winning model to the three other models. These tests revealed that for correct decisions, model 2 (i.e., Confidence ∼ RT*Age + (RT | Participant)), which included the interaction between RT and age, fitted the data best. It was superior to model 1 (the null model), which included only the fixed effect of age [χ^2^(2) = 38.15, *p* < .001], and model 3, which included only the interaction between PF and age [χ^2^(2) = 33.71, *p* < .001]. Moreover, model 4, which included the full interaction between PF, RT and age, did not improve the fit further [χ^2^(4) = 8.64, *p* = .071].

The best fitting model showed significant negative effects of age [β = −0.147, *SE* = 0.039, *z* = −3.75, *p* < .001] and RT [β = −0.109, *SE* = 0.018, *z* = −6.13, *p* < .001] on confidence and a significant interaction between the two factors [β = −0.050, *SE* = 0.017, *z* = −2.96, *p* = .003; Table 1]. Given that we found a significant interaction between RT and age, we computed simple slopes for three values of age (the mean and one SD above and below the mean). The analysis revealed that the negative effect of RT on confidence (i.e., higher confidence for faster responses) increased with older age [Figure 2A; −1SD (i.e., younger adults): β = −0.097, *SE* = 0.026, *z* = −3.75, *p* = .001; mean (middle-aged adults): β = −0.147, *SE* = 0.018, *z* = −8.32, *p* < .001; +1SD (i.e., older adults): β = −0.197, *SE* = 0.023, *z* = −8.53, *p* < .001]. The Johnson-Neyman technique revealed that the effect of RT on confidence became significant from around 24 years of age onwards (higher bound of insignificant interaction effect: 24.41; Figure 2A).

**Table 1.** Regression coefficients (Estimate), standard errors (SE), and associated z- and p-values from the winning generalised linear (beta distribution) mixed-effects model for predicting confidence in correct responses.

| Fixed effects | Estimate | SE | z | p |
| --- | --- | --- | --- | --- |
| (Intercept) | 2.630 | 0.044 | 59.75 | <.001 |
| RT | -0.109 | 0.018 | -6.13 | <.001 |
| Age | -0.147 | 0.039 | -3.75 | <.001 |
| RT*Age | -0.050 | 0.017 | -2.96 | .003 |

**Figure 2.**
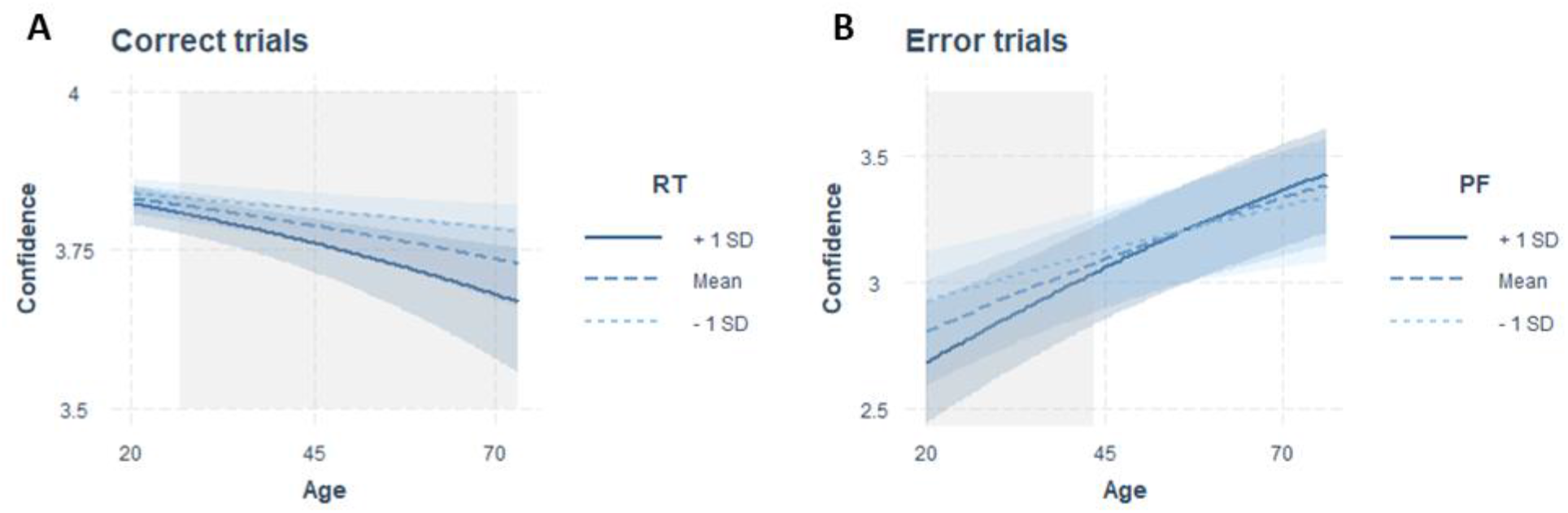
Interaction plots including the predictors of the models predicting confidence best for errors and correct responses. (A) Regression of age on confidence in correct trials with the moderator RT. (B) Regression of age on confidence in error trials with the moderator PF. Regressions are shown for the moderator fixed on the mean (dashed line) and one standard deviation above (solid line) and below (dotted line) the mean. Blue shaded areas indicate confidence intervals, and grey shaded areas indicate the age range in which a significant effect of RT (in correct trials) or PF (in error trials) on confidence is observed, resulting in the significant interaction effect.

#### 3.3.2. Confidence in erroneous decisions

For errors, the best fitting model was model 3 (i.e., Confidence ∼ PF*Age + (RT | Participant)), which included the interaction between PF and age. The likelihood ratio tests revealed that this model fitted the data better than model 1 (the null model) [χ^2^(2) = 8.38, *p* = .015] and model 2, which included the interaction between RT and age [χ^2^(2) = 5.36, *p* < .001], and model 4, which included the interaction between all three factors, did not show an improved model fit, either [χ^2^(4) = 4.47, *p* = .347].

The winning model showed a significant effect of age [β = 0.288, *SE* = 0.066, *z* = 4.40, *p* < .001] and a significant interaction between PF and age [β = 0.080, *SE* = 0.030, *z* = 2.67, *p* = .008], but no main effect of PF [β = −0.018, *SE* = 0.029, *z* = −0.65, *p* = .519; Table 2]. To further unpack the interaction effect, we ran a simple slope analysis. This analysis showed a negative relationship between confidence and PF only for younger adults, while with increasing age, the slope was not significantly different from zero [Figure 2B: −1SD (i.e., younger adults): β = −0.098, *SE* = 0.038, *z* = −2.60, *p* = .009; mean (middle-aged adults): β = −0.018, *SE* = 0.029, *z* = −0.645, *p* = .519; +1SD (older adults): β = 0.061, *SE* = 0.045, *z* = 1.37, *p* = .170]. Computation of the Johnson-Neyman interval showed that above an age of about 44 years (lower bound of significant interaction effect: 43.501), PF was no longer significantly associated with confidence (Figure 2B).

**Table 2.** Regression coefficients (Estimate), standard errors (SE), and associated z- and p-values from the winning generalised linear (beta distribution) mixed-effects model for predicting confidence in incorrect responses.

| Fixed effects | Estimate | SE | z | p |
| --- | --- | --- | --- | --- |
| (Intercept) | -0.085 | 0.065 | -1.31 | .192 |
| PF | -0.018 | 0.029 | -0.65 | .519 |
| Age | 0.288 | 0.066 | 4.40 | <.001 |
| PF*Age | 0.080 | 0.030 | 2.67 | .008 |

## 4. Discussion

This study investigated age-related changes in the relationship between the temporal and motor response parameters RT and PF with decision confidence. Overall, higher confidence was related to faster and less forceful responses. We could further show that, across the entire sample, confidence was associated with both parameters, and these effects were moderated by performance accuracy: While RT was related to confidence in correct responses, peak force was related to confidence in error trials. Finally, age interacted with the response parameters so that with higher age, the effect of RT on confidence was more pronounced, while the effect of PF on confidence was diminished.

We will first focus on the observed age-related associations between confidence and response parameters and subsequently discuss possible interpretations within two different theoretical frameworks.

### 4.1. Behavioural correlates of confidence

In a complex conflict task, we replicated one of the most robust findings on decision confidence, namely a negative relationship between RT and confidence (Fleming et al., 2010; Rahnev et al., 2020). In correct trials, the higher participants rated their confidence in a decision, the faster they had made the decision. In addition, we found a negative relationship between PF and confidence for errors (higher PF was related to lower confidence for the younger participants), which has not been reported before. Observing the latter association is interesting *per se* because participants’ attention was not directed to the applied force in any way (i.e., participants were not aware of the PF assessment), whilst the relevance of speed had been stressed in the instructions. A recent study (Turner et al., 2021a) showed, in a sample of young participants, that when higher levels of force had to be produced to report the (correct) decision, participants’ confidence ratings were higher. While these results do not mirror ours, it should be noted that these findings also cannot be directly compared as their study was conceptually different to our study design and explicitly required participants to produce different force ranges. However, these studies together highlight the added value of assessing response force. In our study, the differential effect of RT and PF for correct and error trials, respectively, further highlights that these are dissociable parameters of a response, supporting a model of Ulrich and Wing (1991; see Armbrecht et al., 2013; Jaśkowski et al., 2000) that RT and RF do not reflect just two sides of the same coin. Further, our findings stress the significance of including incorrect responses as a distinct response type in the corresponding analyses.

Interestingly, the observed associations between the response parameters and confidence differed across the studied age range. While age-related changes in metacognitive performance, and error detection in particular, have been shown across tasks and domains (Harty et al., 2013; Niessen et al., 2017; Palmer et al., 2014), specific characteristics of confidence judgements have rarely been investigated in the context of healthy ageing. The current study revealed a stronger association with increasing age between confidence and RT in correct trials and a weaker association between confidence and PF in errors. Since the current sample of participants covered a broad age range, this constitutes a further step in identifying and understanding age-related changes in metacognitive performance.

We will present two complementary but not exclusive interpretations of the observed agerelated variations in the following. In the first part, we attempt to explain our findings under the assumption that response characteristics simply co-occur with the build-up of confidence. In contrast, in the second part, we assume that response parameters comprise additional information about the decision accuracy that is integrated into confidence during its formation process.

### 4.2. Response parameters as the expression of confidence

One possible framework for explaining the experimental findings is to assume that the level of decision confidence is expressed in the RT or PF of the response indicating this decision, either because confidence defines the response parameters or because a common process drives both confidence and the two parameters. In other words, if a participant is highly confident in a decision, this will affect the speed and the force with which they report this decision. Research has identified multiple stimulus-related characteristics that alter the accuracy of confidence judgements, like relative and absolute evidence strength (Ko et al., 2022; Peters et al., 2017) or evidence reliability (Boldt et al., 2017). If sensory evidence is unambiguous, an easy decision will accordingly lead to high certainty of having made a correct response. In turn, if the participant nevertheless responds incorrectly but changes their mind and detects this error, the certainty of having made an error will be high (i.e., resulting in a low confidence rating). It is intuitive to imagine that high *certainty* of having made a correct or incorrect response (which is identical to very high or very low confidence, respectively) will lead to fast and more forceful responses.

Notably, neither RT nor PF showed the expected pattern of change as would be expected if one or both parameters simply mirrored a decline in confidence with age. Arguably, it might still be possible to explain the differential interactions with age by assuming that multiple other sources (e.g., perception, attention, response selection, motor processes) cause the observed relationships between confidence and the two response parameters. If ageing impacts (some of) these sources differentially, this might result in altered associations between confidence, RT and PF, as observed here. For instance, a cognitive process that is differentially susceptible in older compared to younger adults might affect the RT-confidence relationship but spare the relationship between PF and confidence. However, as we did not systematically investigate these other processes in the present study, we can neither support nor rule out these assumptions.

### 4.3. Modulation of confidence by response parameters

Alternatively, our findings could also be interpreted in line with recent studies postulating that parameters of a response indicating a decision may serve as an additional source of evidence that is integrated into confidence judgements about this decision – especially in ambiguous situations (Filevich et al., 2020; Gajdos et al., 2019; Pereira et al., 2020; Turner et al., 2021a; Wokke et al., 2020). These studies showed that confidence could be altered, for instance, by applying TMS to the dorsal premotor cortex or by instructing participants to move faster (Fleming et al., 2015; Palser et al., 2018). Using very different methodological approaches, these studies mutually indicate that the post-decisional evidence accumulation might incorporate response characteristics of the initial decision into the subsequent confidence rating. Although our study design assessing the relationship of confidence with RT and naturally occurring PF precludes any conclusions regarding the causal direction of effects, it is nevertheless interesting to reflect on our results within this framework.

Looking at the overall relationship between confidence and the two response parameters, our differential findings for errors and correct responses suggest serial processing. First, the RT-related information might be ‘read out’ by the monitoring system and serve as an interoceptive cue about the difficulty of a decision. This assumption is in line with previous work (Dotan et al., 2018; Fleming et al., 2010; Gajdos et al., 2019; Kiani et al., 2014; Rahnev et al., 2020). This interpretation would suggest that the decision-makers arrive at a higher confidence judgement because they also register having responded faster (e.g., via the efference copy (Latash, 2021) or the later representation of their action).

However, this proposed mechanism might exclusively operate in correct trials to refine confidence judgements. For the relationship between confidence and RT in error trials, which were on average slower than correct trials, it must be considered that a variety of aspects can cause errors (e.g., lack of attention, perceptual lapse), and the response profiles of errors are similarly heterogeneous. Therefore, in case of conflict (which is present in error trials), RT might no longer yield reliable information about the task requirements, and the monitoring system might probe PF instead as an alternative response parameter to compensate for the lack of reliable RT when computing confidence. In support, recent work indicated that within a similar speed range for responses, the PF in error trials was related to decision confidence (Stahl et al., 2020). Hence, while RT might not differentiate confidence levels in errors, variations in PF may well capture this information and could therefore be integrated into the final confidence judgement.

Given the frequently described decline of metacognitive abilities with older age, which was also shown in our previous analysis of the current data set (by using the *Phi* correlation coefficient (Nelson, 1984) for the analysis of metacognitive accuracy (Overhoff et al., 2021)), it seems likely that the older adults were lacking relevant input for the computation of confidence, making it harder for them to accurately rate their decisions. Consequently, one possibility is that the stronger association between confidence and RT in older adults might reflect a compensation mechanism. To explain, while our study does not allow for firm conclusions as to why metacognitive accuracy declined in older adults, it appears that the input to the performance monitoring system was diminished (or not adequate anymore) and did not allow for computing confidence with the same level of accuracy as in younger adults. Therefore, it is possible that the stronger reliance on RT (in correct trials) might reflect the attempt to compensate for this by relying more on other sources of input, like the interoceptive feedback about the response speed (Fleming et al., 2010; Palser et al., 2018). However, it remains unclear whether this compensation fails, as, despite more substantial reliance on RT information, confidence judgements were still poorer compared to younger adults. This could be plausible, for example, if the monitoring of response parameters itself might also become poorer with increasing age. Alternatively, it is also possible that this compensation was indeed (somewhat) successful, and without incorporating RT information more strongly, confidence judgements would be even worse. Ultimately, our study cannot resolve this question.

For error trials, we observed that the relationship between confidence and response force diminished with age. One explanation might be related to the finding that healthy ageing has been associated with diminished neural specificity for errors (Endrass et al., 2012; Harty et al., 2017; Overhoff et al., 2021; Park et al., 2010), meaning that older adults might have generally been worse at detecting the errors in the first place. A recent fMRI study has extended these findings by showing that the activity related to error awareness was specifically reduced in older adults (Sim et al., 2020). Based on these findings, our results could be interpreted as another instance of an agerelated error-specific processing deficit. This functional processing deficit might also extend to the sensorimotor feedback of the produced force. The read-out of the response force – which might be used to infer confidence in case of errors – might thus not be readily accessible by older adults and potentially contribute to the demonstrated deficits in metacognitive accuracy.

### 4.4. Limitations

While the simultaneous recording of two response parameters for each response constitutes a strength of the present study, treating RT and PF as equivalent may be problematic. We have discussed RT and PF as separate but comparable features of motor activity, even though their apparent relevance differed largely. Force was produced without constraints, while the time to report the decision was limited to 1,200 ms and exerted considerable time pressure on the participants. Therefore, it would be interesting to examine changes in the modulation of confidence by RT and PF without limiting the time to respond. Moreover, since the RT in a given trial represents the sum of the time for stimulus-related processes (between stimulus onset and the start of the response movement) and the time for motor-related processes (e.g., movement time – the time between starting and terminating a response movement), future studies should additionally assess movement time and its relation to confidence.

As mentioned above, this study cannot resolve the question of causality of the observed associations. Based on the described literature, it is reasonable to speculate that our findings can be explained within the framework of response dynamics informing confidence judgements. However, we have carefully outlined an alternative explanation and acknowledge that both lines of interpretation may be valid in part.

## 5. Conclusion

Corroborating recent evidence, we revealed significant associations between decision confidence and the parameters of the responses indicating this decision. Furthermore, we extended these findings by showing that confidence was associated with fine-grained changes in the time taken to report a decision and the force invested in this response. These relationships were moderated by the accuracy of the response, and, most importantly, changed markedly across the adult life span. This notion should encourage the recording of response force in behavioural experiments whenever possible, as it might uncover specific effects that cannot be revealed by measuring other response parameters, like response times. While a causal explanation of these findings was beyond the scope of this study, one possible interpretation is that the observed agerelated changes in the pattern of associations reflect a mechanism in the computation of confidence and may even constitute one aspect of the frequently observed decline in metacognitive ability with older age.

## Acknowledgements

We thank all colleagues from the Institute of Neuroscience and Medicine (INM-3), Cognitive Neuroscience, the Department of Individual Differences and Psychological Assessment, and the Decision Neuroscience Lab at the Melbourne School of Psychological Sciences for valuable discussions and their support.

