## Supplementary material for "The relationship between response dynamics and the formation of confidence varies across the lifespan"

### Supplement

#### S1. Figures control analyses

In order to get an overview of the distribution of the behavioural parameters of interest independent of confidence, we compared confidence ratings, RT, and PF between errors and correct responses. We assessed the effect of age on the movement parameters RT and PF. Results of *t*-tests and simple linear regressions are reported in the manuscript. Here, we are additionally providing the respective plots for illustration purposes.

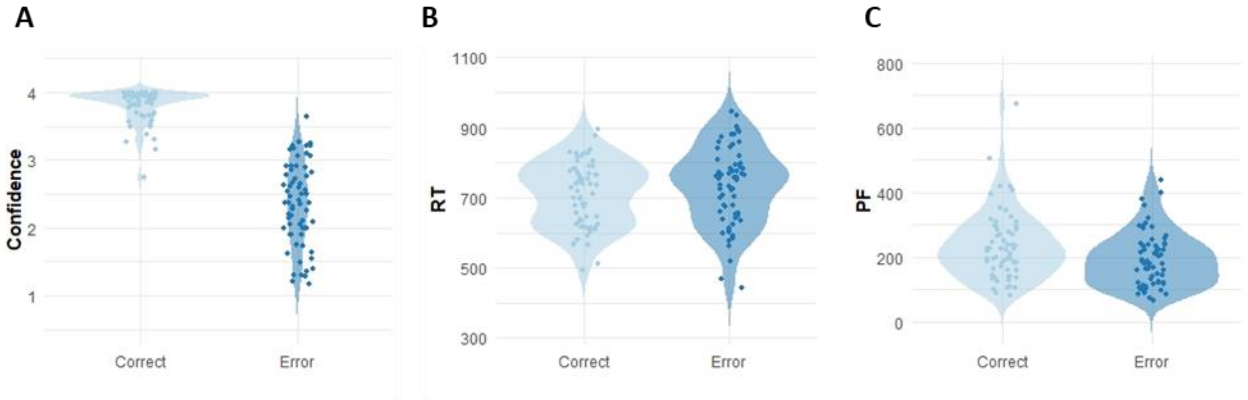


**Supplementary Figure S1.** Distribution of (A) confidence ratings, (B) RT, and (C) PF for errors and correct responses. Dots indicate individual means (for confidence) or medians (for RT and PF), respectively.

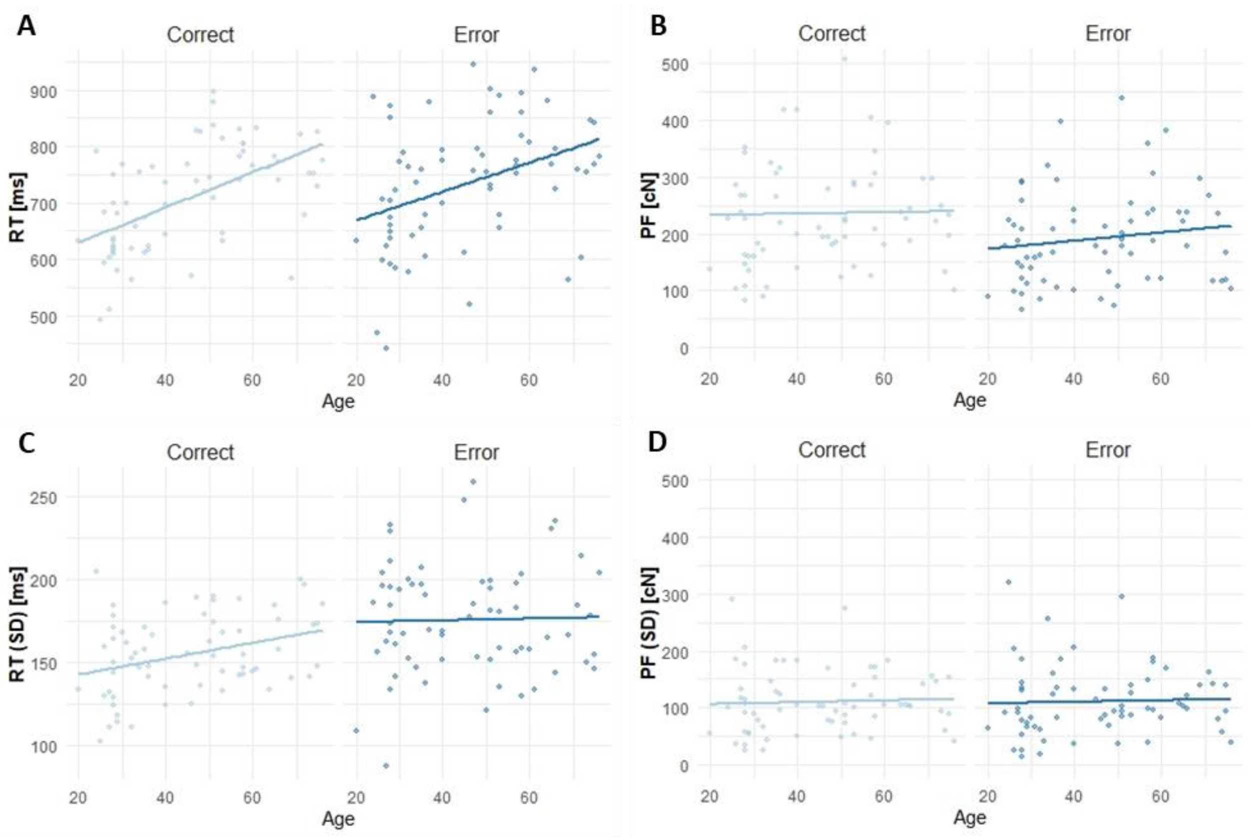


**Supplementary Figure S2.** Regression of RT (left column) and PF (right column) on age for correct and error trials, respectively. Dots and fitted lines in the upper row indicate individual median RT/PF, and dots and fitted lines in the lower row indicate individual standard deviation (*SD*) for RT/PF. Median RT of correct and error trials and *SD* RT of correct trials increase significantly with age.
